# Neural underpinnnings of auditory salience natural soundscapes

**DOI:** 10.1101/376525

**Authors:** Nicholas Huang, Mounya Elhilali

## Abstract

Salience is the mechanism whereby attention is automatically directed towards critical stimuli. Measuring the salience of a stimulus using behavioral methods risks confounds with top-down attention, particularly in the case of natural soundscapes. A distraction paradigm is employed here to measure physiological effects of salient auditory stimuli using electroencephalography. Several such effects are presented. In particular, a stimulus entrainment response is reduced by the presentation of distractor salient sounds. A reduction in oscillatory neural responses in the gamma frequency band is also observed following salient stimuli. These measures are used to identify salient portions of the natural scene. Finally, envelope decoding methods also indicate that salient stimuli attract attention away from other, task-related sounds.

---

Salience is the natural propensity of a stimulus to draw our attention towards it. A siren is a salient sound used to alert drivers on the street to the presence of an emergency vehicle, despite the presence of more routine sounds that may be competing for attention. Salience is also referred to as ‘bottom-up’ attention because it denotes a property of the sensory signal itself that engages cognitive processes of attention. A salient stimulus possesses sensory attributes that allow it to draw attention in an automatic way, regardless of the observer’s state of the mind or object of attention. As such, it differs from voluntary (‘top-down’) attention because it is involuntary and can compete with goal-directed attention, though the two forms of attention intimately interact to shape perception.

The study of salience in audition has largely relied on artificially-constructed stimuli, such as tone and noise tokens or by piecing together short segments of natural sounds (1–7). Such stimuli provide better control over acoustic parameters of the signal as well as more accurate timing of occurrences of salient events. Still, they fall short of reflecting the natural intricacies of realistic sounds in everyday environments. One study (8) did employ extended natural recordings, but their stimuli were of a single setting (meetings) and were generally very sparse.

In the current work, we employ natural sounds based on a salience database previously developed (9). This dataset offers a variety of content and compositions, as well as a wide selection of sound types including speech, music and nature sounds. It includes samples from everyday situations taken from many databases such as YouTube and Freesound, and contains various settings such as a ballgame at the stadium, a concert in a symphony hall, a dog park, a protest in the streets, and many more. A behavioral judgment of salience was obtained for every moment of each natural scene using a dichotic listening paradigm, whereby subjects continuously reported which scenes attracted their attention. While this approach has its shortcomings (see (9)), it offers an almost instantaneous account of the agreement between listeners of the salience of every event in each scene. Other behavioral accounts of perceptual salience have employed paradigms such as manual labeling (8) or overt behavioral comparisons (4). In all these approaches, by asking listeners to offer their salience judgment either discretely or continuously, these measures are likely to be confounded by top-down attentional effects due to the deliberate and voluntary nature of the responses.

A preferred approach to probing salience is to rely on physiological markers of salience. These measures circumvent the difficulty of separating the effects of bottom-up and top-down attention because they involve no voluntary response. Unfortunately, unlike gaze in visual salience, measuring auditory salience is a particularly challenging task given the lack of obvious physical markers of what/where a person’s auditory attention is located. One physiological marker used in some studies is pupilometry, where recent work has suggested a positive correlation between stimulus salience and pupil size (10, 11). Unfortunately, this connection needs to be further investigated as there is evidence that this correlation may be largely driven by loudness rather than a full account of salience of the acoustic signal; as well as being modulated by local rather longer-term contexts that often guide our judgments of salience (9, 12).

An alternative to pupilometry are direct measures of brain activity. A variety of methods exists for recording responses from the brain. Studies of auditory attention have used blood oxygenation levels recorded using fMRI (13–15), electrical signals measured using Electrocorticography (16, 17) or Electroencephalography (EEG) (18–25), and magnetic signals recorded using Magnetoencephalography (26–28). While this body of existing literature has shown strong modulation of brain activity due to top-down, task-related attention, the present study is conducted to discern whether similar effects occur as a result of bottom-up attention. To this end, EEG has been chosen as the recording modality; EEG benefits from having high temporal resolution, being non-invasive, and being relatively inexpensive compared to other recording methods.

A large number of attentional EEG studies have relied on event related potentials (ERPs), which are time-locked to the onset of the stimulus. (28–34). With continuous natural scenes, such onsets are not available, and other measures are required. Possibilities include continuous neural responses such as stimulus-following responses or steady-state phase-locked entrainment, which have been reported for both stimulus trains (35, 36) and for steady state sounds (37). These responses have been found to be affected by top-down attentional state (38, 39). Changes in oscillatory activity in higher frequency bands such as the gamma band have also been attributed to attentional shifts (17, 40, 41). Variations in both of these signals are examined in this study.

In addition to these metrics, a popular, more recent technique in probing continuous neural responses is envelope decoding. The goal of decoding is to ‘infer’ the stimulus from the neural response, hence reflecting the degree of fidelity of stimulus encoding in brain activation patterns (29–34). Numerous studies have shown that the quality of envelope decoding using both linear and nonlinear decoders is much stronger for attended stimuli as opposed to unattended backgrounds; hence reflecting a strong effect of task-directed attention.

Both ERP and continuous responses are ideal candidates to probe effects of bottom-up attention. The use of natural stimuli in the current study makes both analyses slightly challenging. Measurement of event-related potentials requires precisely-aligned event onsets, which are not feasible for a continuous sound stream. Instead, we present listeners with a concurrent sequence of tones with clear onsets and offsets to allow a time-locked analysis. This tone sequence provides a reference for temporal alignment. In addition, it offers a crucial attentional anchor for the experimental paradigm, since subjects are asked to attend and perform a task relative to the tone sequence; hence ensuring that modulatory effects of salient events from the natural scenes are truly emerging in a bottom-up fashion.

The analysis of continuous responses and decoding measures is well suited to dynamic stimuli such as natural sounds used in the current paradigm. Overall, the current design tries to balance the richness of using a natural stimulus with the controlled structure of a competing task hence regulating the listeners’ attention away from the natural scenes while probing effects of salient events in brain responses to the scene itself as well as the competing tone sequence, that is the object of attention throughout the experiment.

## Results

Human listeners performed an amplitude-modulation detection task by attending to a tone sequence while neural responses using Electroencephalography (EEG) were collected. Concurrently, they were presented with a busy background scene that they were asked to ignore (Fig. 1). The background scenes were taken from the JHU DNSS (Dichotic Natural Salience Soundscapes) Database for which behavioral estimates of salience have been obtained (9). The Huang and Elhilali study identified salient events in each background scene along with their salience strength (see Methods for details).

**Fig. 1.**
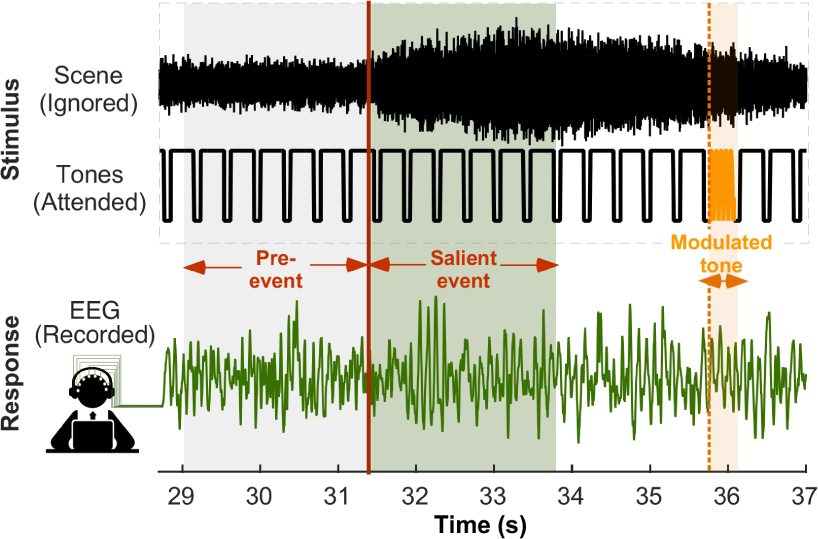
Stimulus paradigm during EEG recording. Two stimuli are presented concurrently in each trial: a periodic tone sequence and a recording of a natural auditory scene. Subjects attend to the tone sequence to perform a modulation detection task. Example stimuli and EEG recordings are shown for a short portion of one scene, along with an example salient event and modulated target tone.

The presence of a regular tone sequence in the stimulus induces a strong phase-locked response to the tone presentation rate at 2.6Hz despite the concurrent presentation of a natural scene. Figure 2A shows the grand average spectral profile of the neural response observed throughout the experiment. The figure clearly displays a strong energy at 2.6Hz, with a left-lateralized fronto-centralized response stereotypical of an auditory cortex source (Fig. 2A, inset). The phase-locked peak at exactly 2.6Hz, normalized by dividing by the mean energy at neighboring frequencies (*±*0.5Hz around 2.6Hz), is significantly greater than one, across all subjects and electrodes [t(1501) = 4.18, p = 3.1*10-5].

**Fig. 2.**
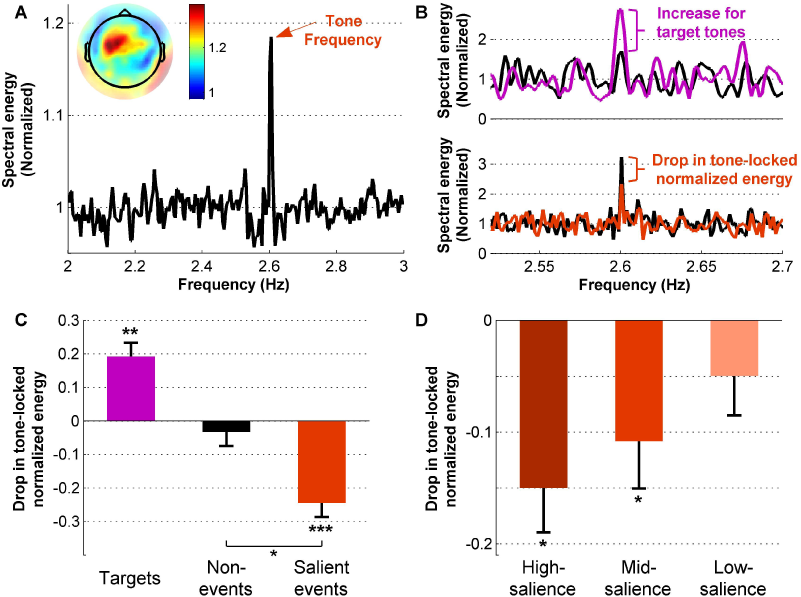
Phase locking results. **(A)** Spectral density across all stimuli. The peak in energy at the tone presentation frequency is marked by a red arrow. Inset shows average normalized tone locking energy for individual electrodes. **(B)** Spectral density around target tones (top) and salient events (bottom). Black lines show energy preceding the target or event, while colored lines depict energy following. **(C)** Change in phase locking energy across target tones, non-events, and salient events. **(D)** Change in tone locking energy across high, mid, and low salience events. Error bars depict *±*1 SEM.

We look closely at modulation of this phase-locked response both by top-down attention in presence of the target tone as well as by bottom-up attention coinciding with the presence of a distracting salient event. Focusing on phase-locking near the target modulated-tone shows an increase in 2.6Hz power relative to the overall level, reflecting an expected increase in neural power induced by top-down attention (Fig. 2B-top). Interestingly, the same phase-locked response is notably reduced when tones coincide with salient events in the background (Fig. 2B-bottom). The spectral profile shows a clear drop in phase-locking at exactly 2.6Hz when the tone co-occurs with a salient event (red curve) indicating a potential marker of distraction effect from the concurrent scene, despite subjects’ attention being directed to the tone sequence.

Separating the phase-locking effect between tones near a the modulated target, salient events and those ‘away’ from targets or salient events shows a clear distinction between the three groups. We quantify this change in phase-locking across all subjects by comparing the neural response during the window of interest (target tone or salient event) relative to a preceding period (Figure 1). We also contrast the same variability in phase-locking at randomly chosen tones away from salient events or targets. Figure 2C shows that phase-locking to 2.6Hz near the modulated target tone increases significantly [t(443) = 4.65, p = 4.43*10^−6^], whereas it decreases significantly following salient events [t(443) = −5.89, p < 10^−7^]. A random sampling of portions of the scene away from salient events does not show any change [t(443) = −0.78, p = 0.43].

The top-down attentional effect due to tone targets is significantly different than inherent variability in phase-locked response when compared to non-events [t-test, t(886) = 3.81, p = 1.48*10^−4^]; while distraction due to salient events induces a decrease in phase-locking that is significantly different than inherent variability in phase-locked response when compared to non-events [t-test, t(886) = −3.58, p = 3.66*10^−3^]. This decrease is modulated in strength with the level of salience of the concurrent events. The decrease is strongest for events with a higher level of salience [t(443) = −3.78, p = 1.8*10^−4^]. It is also significant for events with mid-level salience [t(443) = −2.57, p = 0.01] with no significant reduction in tone locking for events with the lowest salience [t(359) = 1.33, p = 0.20] (Fig. 2D).

Next, we probe other markers of attentional shift and focus particularly on the Gamma band energy in the spectral response (17). We analyze the spectral energy across the frequency spectrum in a time period surrounding target tones and salient events. Figure 3A depicts a time-frequency profile of neural energy before and after modulated target tones (0 on the x-axis denotes the start of the target tone). A strong increase in Gamma activity occurs after the onset of the target tone. Figure 3B shows the same time-frequency profile of neural energy but now relative to salient events. The figure clearly shows a decrease in spectral power around the high Gamma range between 70-120Hz following salient events.

**Fig. 3.**
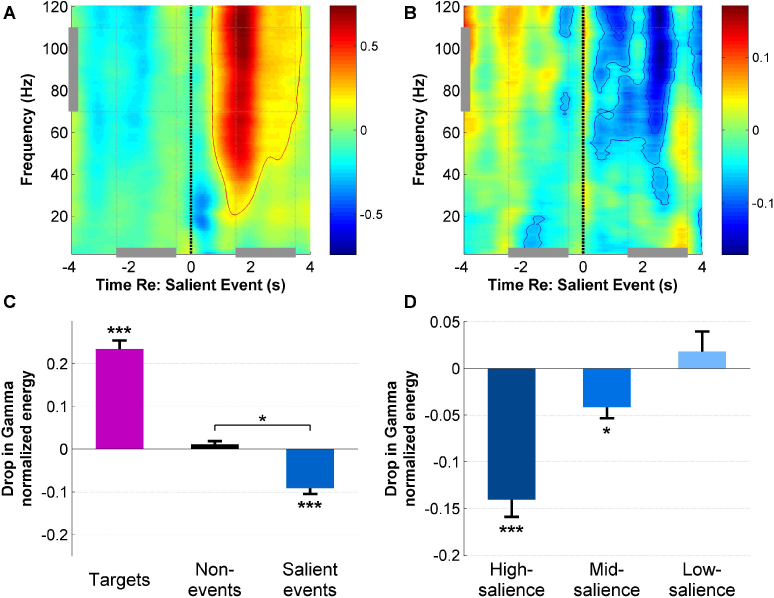
High gamma band energy results. **(A)** Time frequency spectrogram of EEG recordings around modulated target tones, averaged across central and frontal electrodes. **(B)** Time frequency spectrogram around salient events. **(C)** Change in energy in the high gamma frequency band (70-110 Hz) across target tones, non-events, and salient events. **(D)** Change in high gamma band energy across high, mid, and low salience events. Error bars depict *±*1 SEM.

Figure 3B quantifies the variations of Gamma energy relative to targets, salient events and non-event control tones. Gamma band energy increases significantly following the target tones for all subjects [t(443) = 11.5, p < 10^−7^]. It drops across salient events [t(443) = −6.83, p < 10^−7^] compared to control non-event segments which show no significant variations in high-Gamma energy [t(443) = 1.5, p = 0.13]. The increase in spectral energy around the Gamma band is significantly significant in a direct comparison between target and control tones [t(886) = 10.3, p < 10^−7^]. Similarly, the decrease in spectral energy around the Gamma band is significantly significant when comparing salient events against control non-events [t(886) = 6.68, p < 10^−7^]. As with the decrease in tone locking, the Gamma band energy drop is more prominent for higher salience events [t(443) = −7.72, p < 10^−7^], is lower but still significant for mid-level salience events [t(443) = −3.64, p = 3.02*10^−4^] but not significant for low-salience events [t(443) = 0.84, p = 0.40] (Fig. 3C).

Next, We focus on the predictive power of neural markers of auditory salience and specifically examine how predictive are tone-locking and Gamma energy about the alignment of each tone in the sequence with a salient or non-salient events in the background scene. We train a classifier to label each tone as attended (i.e. coinciding with no salient event) vs. distracted (i.e. coinciding with a salient event) and probed prediction accuracy with each neural marker. Figure 5 shows that classification accuracy, measured by the area under the ROC curve, is at chance level when using variations in Gamma-band energy [t(9) = −0.10, p = 0.92]. However, tone-locking energy yield significant predictions above chance [t(9) = 3.24, p = 0.01]. The best accuracy is achieved when including both tone-locking and Gamma energy [t(9) = 4.50, p = 0.0015], suggesting that Gamma-band energy does contribute complementary information that is not above the noise-floor on its own but is nonetheless informative about presence of distractor salient events in the background.

**Fig. 4.**
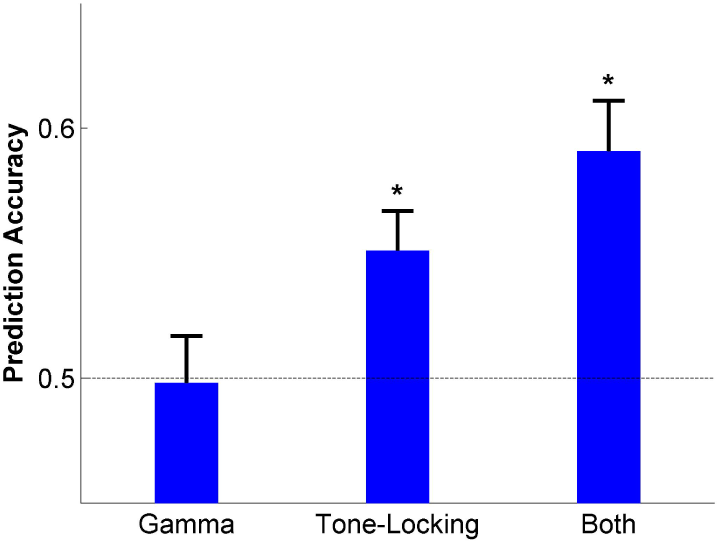
Event Prediction Accuracy. Average event prediction accuracy (area under the ROC curve) using only high gamma band energy, only tone-locking energy, and both features. Error bars depict *±*1 SEM.

**Fig. 5.**
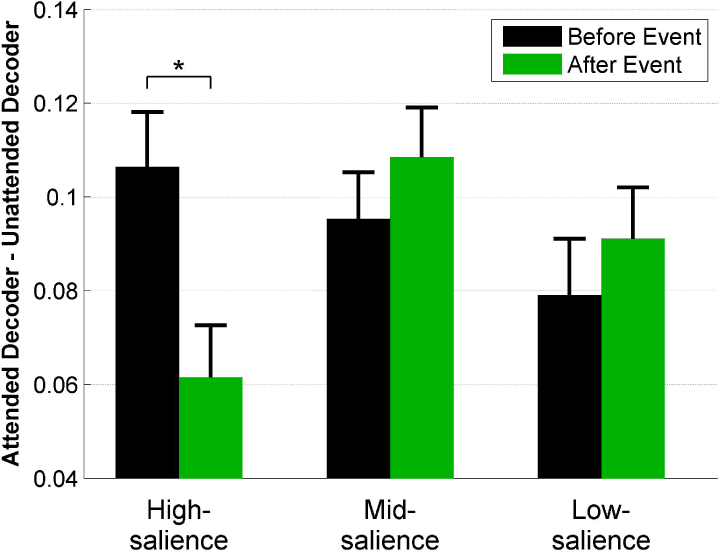
Envelope Decoding Performance. Envelope decoding performance of the attended stimulus (tone sequence) minus the unattended stimulus (natural scene). Increases indicate relatively higher attention to the tones, decreases indicate relatively higher attention to the scene. Decoding performance shown before and after high, mid, and low salience events. Error bars depict *±*1 SEM.

With the use of dynamic natural scenes in the background, it is possible to further explore disruptions to the fidelity of the neural encoding of the stimuli by examining the accuracy of envelope decoding before and after salient events. Figure 5 reports the change in decoding accuracy after salient events grouped by salience strength. The envelope decoding accuracy for the tone sequence drops in comparison to decoding performance for the natural scene following events of high salience [t(204) = −2.77, p = 0.0061]. This decrease is not observed for events of a lower salience level [Mid: t(226) = 0.90, p = 0.37; Low: t(214) = −1.91, p = 0.06].

## Discussion

Attention modulates brain activity, and bottom-up auditory attention is no exception. A number of feature have been identified that can be used to identify salient portions of an auditory scene, using a distraction paradigm. These features include spectral features (locking to the presentation frequency of the attended tone sequence and energy in the high gamma band) and temporal features. Using these features and a classification method, salient events can be identified within these scenes with an accuracy of around 60%, as measured by the area under the ROC curve. In addition, the ability to decode the envelope of the stimulus relative to the envelope of the task-related tone sequence also improved after salient events. However, this increase was only observed after the events with the highest salience values.

The observed drop in tone-locking after salient events directly indicates that subjects were less able to follow the tone sequence during those times. This effect was particularly pronounced for higher salience events. In contrast, energy at the tone presentation frequency increased significantly upon presentation of the modulated target tone, providing further evidence that entrainment to the tone sequence serves as a measure of attention to the task.

Unfortunately, in order to obtain uncontaminated EEG data, target tones were not placed in the post-event region. As such, the behavioral data cannot corroborate this statement. Another limitation of this analysis is that it is difficult to directly compare the magnitude of change between targets tones and salient events. Within the stimulus paradigm, there were far more events (234) than modulated target tones (79). Because windows around targets/events were directly concatenated, the power spectral density of the salient events had greater frequency resolution. Thus, energy at the tone presentation frequency was better isolated from neighboring frequencies for salient events (Fig. 2B).

A shift in attention after salient events occur is also indicated by the drop in high gamma band activity. Gamma band activity is associated with increased task attendance (Gruber et al., 1999), and indeed energy at these higher frequency bands does increase following the modulated target tones. Thus, as with the decrease in tone-locking, this effect indicates a decrease in task attentiveness of the subjects immediately following salient events. Again, in parallel with the tone-locking results, this effect is greatest for the highest salience events.

The event prediction performance is predominantly led by the tone locking activity, which is more consistent when looking across all events compared to the the gamma band energy. However, although prediction performance is poor when using only activity in the gamma band, including it as additional information with the tone locking activity does improve the quality of the prediction.

Envelope decoding performance for the natural scene was relatively low compared to the decoding of the attended tone sequence. In addition to general noise from EEG recording and the simplicity of the tone envelope, this lower performance is at least partially the result of the subjects attending to the tone sequence rather than to the natural scene. However, the decoding performance did show a relative increase for the scene compared to the tone sequence after high salience events. This change suggests that these strong events were able to at least partially attract the subjects’ attention away from their task stimuli for a brief time.

One interpretation of the results is that the EEG is only recording responses to basic acoustic features rather than salience. For instance, the scenes may be louder immediately following all acoustic events than they are at any other time. Although there is a small difference in the average level of some acoustic features directly following salient events, however, this difference is highly inconsistent. The distribution of the levels of these features, when comparing two seconds after events to the rest of the scene, is highly overlapping. This quality is shown using several different measures in Table 1.

**Table 1.**
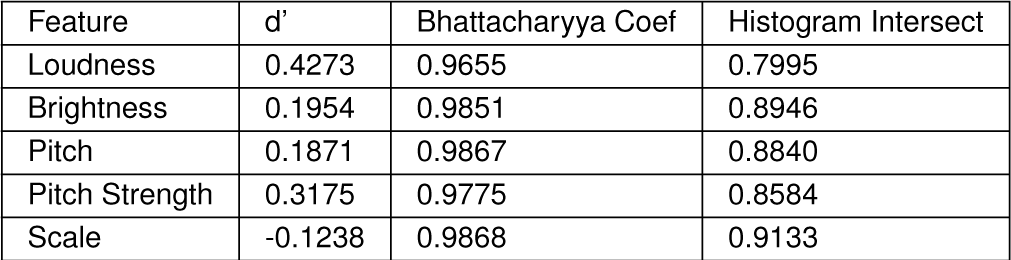
Comparison of the distribution of several acoustic features immediately after events and across the rest of the scene.

In this study, distraction has been used as a method for measuring a subject’s attention. A salient stimulus can draw attention away from an unrelated task, lowering performance on that task. Such a change in performance can be measured both by physiological measurements as it was in this study, or by changes in behavioral response. The target stimuli and events were well-separated in order to avoid confounding their effects; as a result, however, effects of the distractors on the behavioral response could not be measured. One natural following step would be to present a more continuous behavioral task concurrently with a natural scene, and then to examine effects of salient events on task performance.

## Materials and Methods

### Stimuli

Auditory stimuli consisted of scenes from a previous study of auditory salience in natural scenes (9). This JHU DNSS (Dichotic Natural Salience Soundscapes) database consists of twenty natural scenes, each roughly two minutes in length. These scenes were drawn from many sources (including Youtube, Freesound, and the BBC Sound Effects Library), and encompassed a wide range of settings. Some scenes were acoustically sparse, such as bowling sounds in an otherwise quiet bowling alley. Others were acoustically dense, such as an orchestra playing in a concert hall or a busy cafeteria.

Salience within these scenes was measured using a behavioral paradigm based on dichotic listening. Human listeners were presented with two simultaneous scenes and asked to indicate which scene they were attending to in a continuous fashion. The proportion of subjects attending to a scene compared to all other scenes was defined as salience. Peaks in the derivative of this salience measure were defined as salient events. The strength of salient events was further defined as a linear combination of this slope of salience and the maximum value of salience in a short window (four seconds) after the event. This salience strength was used to rank the events as high-salience, mid-salience, or low-salience.

In the current study, each scene was presented individually, one at a time concurrently with a 440 Hz tone sequence, forming a single trial (Figure 1). Tones were presented at a presentation rate of 2.6 Hz and had a duration of 300 ms, with 10 ms cosine on and off ramps. Within each trial, three to five of these tones were amplitude modulated at 64 Hz with a modulation depth of 0 dB. These amplitude-modulated tones were constrained to be at least 8 tone periods away from any salient event within the concurrent natural scene. The stimuli (concurrent scene and tone sequence) were presented binaurally to both ears via a pair of ER-3A insert earphones.

### Participants and Procedure

Twelve subjects (ages 18-28, nine female) with no reported history of hearing problems participated in this study. All experimental procedures were approved by the Johns Hopkins University Homewood Institutional Review Board (IRB), and subjects were compensated for their participation. Subjects were tasked with counting the number of amplitude modulated tones within each scene, and they reported this value after each scene ended using a keyboard. Each subject heard each of the twenty scenes one time only. They were instructed to focus on the tone sequence and to treat the auditory scene as background noise.

### Electroencephalography

EEG measurements were obtained using a 128-electrode Biosemi Active Two system (Biosemi Inc., The Netherlands). Reference electrodes were placed at both left and right mastoids; and electrodes were also placed below and lateral to each eye, in order to monitor eye blinks and eye movements. Data were initially sampled at 2048 Hz.

24 electrodes surrounding and including the Cz electrode were considered to be in the ‘Central’ area, while 22 electrodes surrounding and including the Fz electrode were considered to be in the ‘Frontal’ area (24). Eight electrodes were located in the intersection of these two sets; unless otherwise stated, statistical tests were performed using the union of the Central and Frontal electrodes.

#### Preprocessing

EEG data were analyzed using MATLAB (Mathworks Inc., MA), with both FieldTrip (42) and EEGLab (43) analysis tools. The data were first demeaned and detrended, then a highpass filtered was applied (3rd order Butterworth filter with 0.5 Hz cutoff frequency). The data were then resampled to 256 Hz. Power line noise at 60 Hz was then removed using the Cleanline plugin for EEGLab (44). Outlier electrodes were removed using a joint improbability criterion on energy at 20-40 Hz, with a threshold of 2.5 standard deviations. This method excludes electrodes that exceed the average energy in that frequency band across all electrodes by more than 2.5 standard deviations. The data were then re-referenced using a common average reference.

#### Data Analyses

For event-based analyses (tone locking and gamma band energy), EEG data were divided into epochs centered on the onset of the tone that was closest in time to each salient event (as defined in Stimuli) or centered on the onset of the modulated target tone. Each of these ‘epochs’ was ten seconds in duration. Noisy epochs were excluded using a joint probability criterion on the amplitude of the EEG data, which rejects trials with improbably high electrode amplitude. A local (single electrode) threshold of 6 standard deviations away from the mean was used, as well as a global (all electrodes) threshold of 2 standard deviations. The data were then decomposed using Independent Component Analysis (ICA). Components stereotypical of eyeblink artifacts were manually identified and removed by an experimenter.

Tone-locking analysis was performed by taking the Fourier Transform of the data, using windows of approximately 2.3 seconds (6 tone periods). Windows corresponding to all events of a given category were first concatenated together before the Fourier Transform was taken. This concatenation was utilized because of the low frequency of interest coupled with the short stimulus duration, and to achieve higher frequency resolution. The concatenated signal was zero padded to a length corresponding to 260 event windows in order to maintain the same frequency resolution for all conditions. The energy at 2.6 Hz was normalized by dividing by the average energy at surrounding frequencies (between 2.55 and 2.65 Hz). For salient events the change in tone locking power was defined as the tone locking power using the first six tones after the events, minus the tone locking power using the six tones prior to the event. For the modulated target tones, this change was defined as the tone locking power of using the target tone and the following five tones, minus the tone locking power using the six tones prior to the target. For illustrative purposes, tone-locking analysis was also performed over the full scene without dividing the data into epochs. (Fig. 2A)., High gamma band analysis was also performed by taking the Fourier Transform of the data in two-second windows around the event or target. The energy at each frequency and each electrode was normalized by dividing by the mean power across the entire event window after averaging across trials, and then converted to decibels (dB). The average power between 70 and 110 Hz was taken as the energy in the high gamma band. The decrease in gamma band power was defined as the high gamma energy in the window containing 1.5 and 3.5 seconds after a salient event minus the high gamma energy between 2.5 and 0.5 seconds before that salient event.

A set of time points that were not near salient events was selected to serve as a control. These control tones were randomly selected from the set of tone onsets, with the criteria that none of these time points were within three seconds of a salient event nor within three seconds of each other. Tone locking and gamma band analyses were repeated using these control periods.

### Predictions

Event prediction was conducted on a per-tone basis. Tones within 5 seconds of an event were included in the prediction. Each tone was categorized as either occurring before the event or after the event. Any tones that were both within 5 seconds before an event and 5 seconds after an event were removed from the analysis. Two features were used in the classification: energy at the tone presentation frequency and gamma band energy. Both measures were averaged across central and frontal electrodes for each of the twelve subjects, resulting in twenty-four features used for classification.

Classification was performed using Support Vector Machine (SVM) (45). For each tone, the classifier was provided feature data from each subject separately. Ten-fold cross-validation was performed by dividing all of the tones from each scene into ten distinct equal portions. During each of ten iterations, one group of tones was used as test data and the remainder used as training data, producing one SVM classification value for each tone.

Once trained, the distance of each tone to the SVM hyperplane was used to rank the tones in order of most to least likely to be immediately following a salient event, since that distance is proportional to the log odds of belonging to a specific class (46). A receiver operating characteristic (ROC) curve was constructed by applying varying thresholds on that ranking, with tones above each threshold predicted as following an event and the remainder predicted as not following an event (47). The area under the ROC curve was then calculated using the trapezoidal method.

### Envelope Decoding

Envelope decoding was performed using a linear decoder as described in O’Sullivan et al. (48). Briefly, decoders were trained to decode the envelope of the stimuli from the EEG data. The attended stimulus (tone sequence) and the unattended stimulus (natural scene) envelopes were decoded separately. The stimulus envelopes were extracted using an 8 Hz low-pass filter followed by a Hilbert transform. The EEG data itself was bandpass filtered between 2-8 Hz (both of these filters were 4^th^ order Butterworth filters). Each linear decoder was trained to reconstruct the corresponding stimulus envelope from the EEG data, using time lags of up to 250 ms on all good electrodes. To avoid overfitting, two decoders (one attended and one unattended) were trained for each of the twenty scenes, using data from the remaining nineteen, resulting in forty decoders per subject.

The quality of these reconstructions was then evaluated by taking the correlation between the original stimulus envelope and the decoded envelope within a sliding window of one second length. Attention to the tone sequence for each window was calculated as the reconstruction quality of the tone sequence minus the reconstruction quality of the background scene. This differential envelope decoding performance before and after salient events was evaluated, with the performance between 0.5 and 1.5 seconds before the event compared to the performance between 0.5 and 1.5 seconds after the event.

